# From manual analysis to automation: scaling passive acoustic monitoring for the endangered Saimaa ringed seals

**DOI:** 10.1101/2024.05.06.592639

**Authors:** Adrià Solana, Mairi Young, Climent Nadeu, Mervi Kunnasranta, Ludwig Houégnigan

## Abstract

Passive acoustic monitoring (PAM) offers a non-invasive method for monitoring elusive pinnipeds, but manual analysis of large recording datasets limits its scalability. For the endangered Saimaa ringed seal (*Pusa saimensis*), PAM provides a rare opportunity to study breeding-season behavior beneath seasonal ice cover. We evaluated automated methods for detecting and characterizing the species’ distinctive knocking vocalizations using recordings from Lake Saimaa. Annotated data (*n* = 12 565 calls) were used to develop pulse repetition rate (PRR) estimation and call-detection systems. The best-performing PRR estimator matched manual measurements with a mean absolute error of 1.50 Hz, while a spectrogram-based convolutional neural network detected knocking calls with a mean F1-score of up to 99.28%. These results show close agreement with manual approaches, indicating that automated detection and characterization are achievable at a standard that could substantially reduce PAM analysis effort. This represents an important step toward scalable, long-term acoustic monitoring of this endangered seal.

## Introduction

The Saimaa ringed seal (*Pusa saimensis)* is an endangered pinniped endemic to the freshwater Lake Saimaa in Eastern Finland. The population was historically considered a game species and actively hunted between 1882 and 1948, before receiving legal protection in 1955 (Kunnasranta et al., 2021). Conservation efforts over recent decades have brought the population from around 100 individuals in 1984 to over 500 today (Kunnasranta et al. 2021; https://www.metsa.fi/suojelu-ja-hoito/lajien-suojelu/saimaannorppa/norppakannan-seuranta/). Nevertheless, this small and fragmented population continues to face significant threats, including bycatch mortality, human disturbance, and climate change, compounded by the low genetic diversity resulting from its long isolation and historical bottleneck (Kunnasranta et al. 2021). Understanding how the population responds to these ongoing and emerging pressures depends on reliable, long-term monitoring capable of tracking the species across its full range and annual cycle.

Passive acoustic monitoring (PAM) provides a non-invasive, cost-effective approach for continuously monitoring wildlife across time and space. Deploying autonomous acoustic recording devices over extended periods generates large amounts of data, which can provide important insights into animal behavior, distribution, abundance, and environmental interactions of various taxa (Gibb et al. 2019; Teixeira et al. 2019). PAM is widely used for monitoring marine mammals, which rely on underwater vocalization for orientation, feeding, mating, nursing, and other social communications (Fleishman et al. 2023). In addition, this method is particularly advantageous for species in habitats where visual observation is challenging, enabling long-term systematic monitoring of vulnerable populations. For the Saimaa ringed seal, this is especially valuable as key reproductive behaviors occur beneath seasonal ice and snow cover, making it the one part of the species’ annual cycle that cannot be directly observed by any other current monitoring method (Sipilä 1990). PAM has already been shown to be successful, with recent work using long-term acoustic data to reveal seasonal patterns in knocking vocalizations that coincide with the presumed mating period, providing the first evidence of the timing of mating in this species (Young et al. 2025). This finding establishes vocal activity as a biologically meaningful indicator of breeding-season behavior and demonstrates PAM’s potential as a valuable complement to current monitoring strategies. Building on this, PAM is now being expanded across a wider network of sites within Lake Saimaa, with the aim of using vocal activity to confirm the presence or absence of seals at the population scale and to monitor reproductively important behavior beneath the ice on an ongoing basis. However, current approaches to analyzing acoustic data rely on manually processing the data, requiring trained analysts to visually and audibly inspect the data to identify target seal calls. Each hydrophone can generate thousands of hours of continuous recordings over a single deployment, and reviewing this volume of audio often requires weeks to months of dedicated analyst effort. This labor-intensive manual process is a current bottleneck limiting the scalability of PAM as a long-term monitoring tool.

Recent advances in machine learning, particularly deep learning, offer a potential solution, providing methods capable of efficiently processing large acoustic datasets (Caruso et al. 2020; Kirsebom et al. 2020; Allen et al. 2021; White et al. 2022; Ziegenhorn et al. 2022; Miller et al. 2023). In particular, convolutional neural networks (CNNs) have proven effective at automatically identifying the sounds of other seal species (Lu et al. 2021; Escobar-Amado et al. 2022), suggesting a similar approach could be applied to Saimaa ringed seals. In this study, we test whether Saimaa ringed seal knocking vocalizations can be reliably detected and characterized automatically, to the standard needed to reduce manual review as the limiting step in acoustic monitoring of this species. Specifically, we assess how accurately a call’s pulse rate -- a key acoustic property of the knocking call that is currently measured manually -- can be estimated automatically. We then test how reliably calls can be distinguished from background noise and other recorded sounds, using several machine learning models. Establishing this is a necessary step toward scaling PAM into a routine, long-term monitoring tool for the Saimaa ringed seal.

## Material and methods

### Acoustic data

This study utilizes acoustic data and annotated knocking calls originally collected and analyzed in Young et al. (2025). Data were collected using four SoundTrap 600 STD hydrophones (Ocean Instruments, New Zealand) deployed in the Haukivesi basin of Lake Saimaa (61°05’ to 62°36’ N, 27°15’ to 30°00’ E), Eastern Finland (Fig. 1). Recordings were made continuously at a sampling rate of 96 kHz and spanned two breeding seasons: a single hydrophone (H_D_) deployed from January to July 2022, and three hydrophones (H_A_, H_B_, and H_C_) deployed between December 2022 and June 2023. Full details of data collection and extraction, including manual annotation and manual measurement of call characteristics, are given in Young et al. (2025).

**Fig 1.**
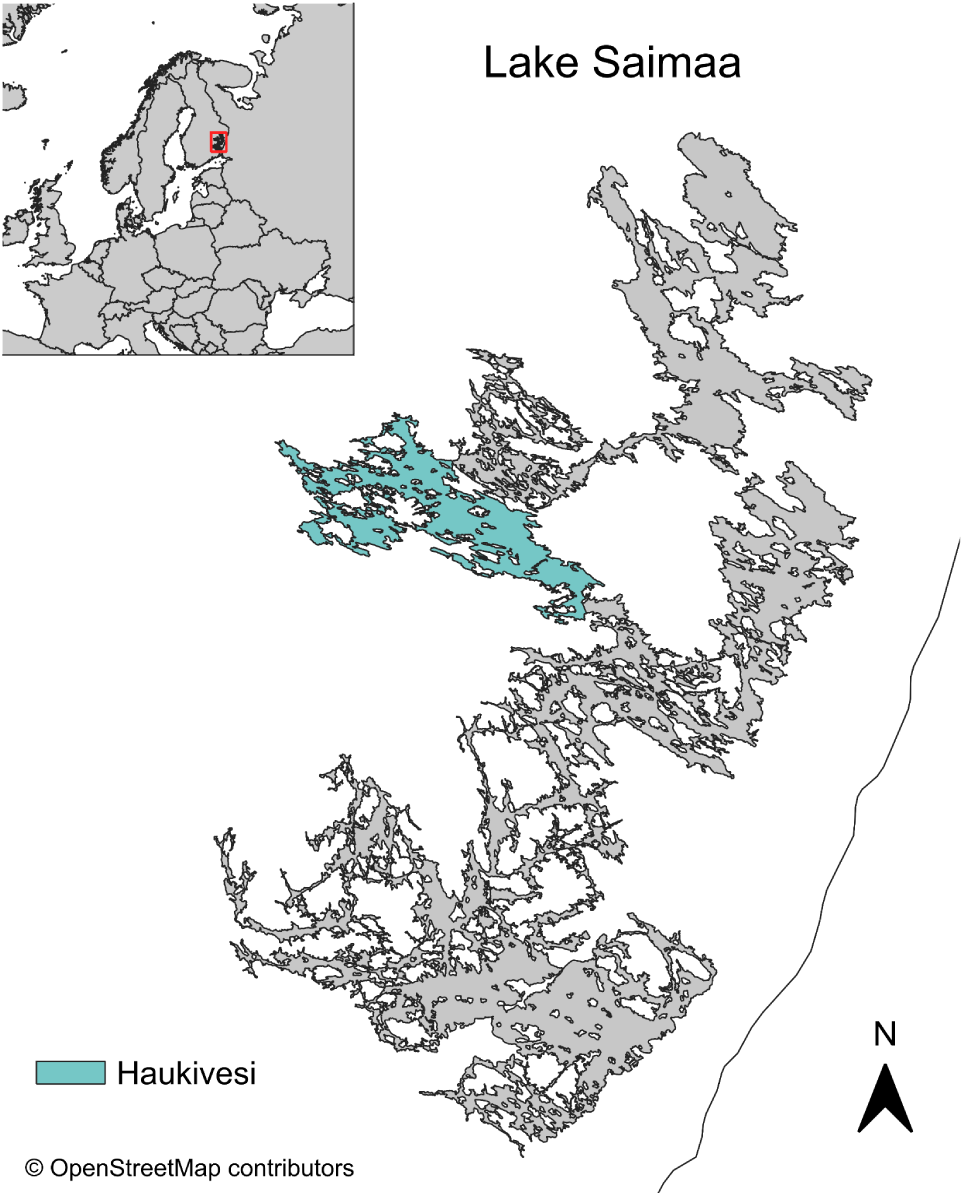
Map of Lake Saimaa with the Haukivesi basin.

Across all recordings, a low-frequency knocking call (Fig. 2) was the only ringed seal vocalization type detected. The call structure was consistent with knocking calls described previously in Saimaa ringed seals and in other ringed seal populations (Kunnasranta et al. 1996; Rautio et al. 2009; Mizuguchi et al. 2016). In this dataset, the call was found to show a seasonal pattern linked to breeding behavior (Young et al. 2025). Being both the sole vocalization type recorded and closely tied to a key reproductive behavior, the knocking call is a natural target for long-term acoustic monitoring. Furthermore, its occurrence across the dataset (2 621 annotated events in H_A_, 24 in H_B_, 953 in H_C_, and 8 967 in H_D_, identified through manual review) provided a substantial foundation on which to develop automated detection and characterization methods.

**Fig 2.**
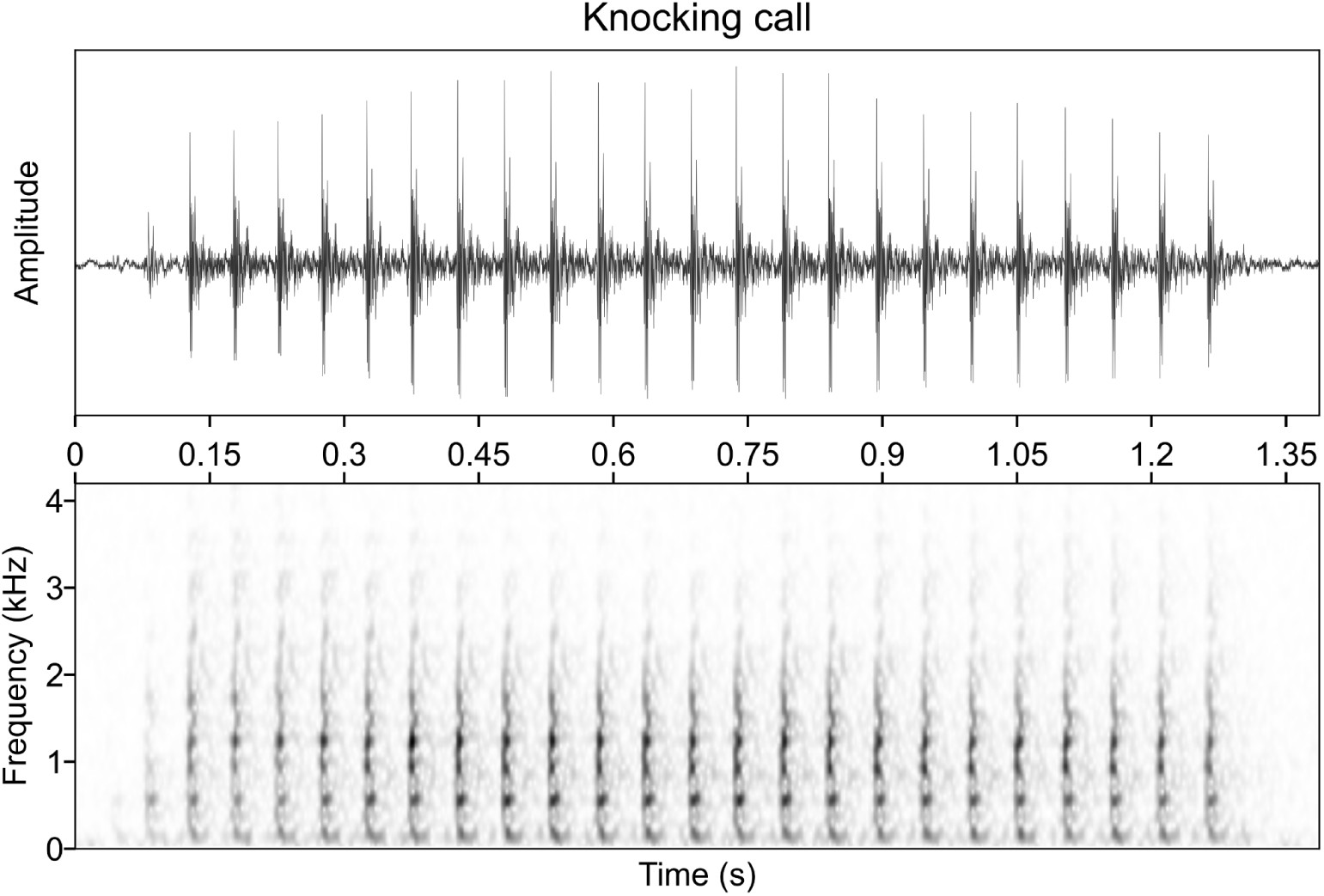
Waveform and spectrogram of a Saimaa ringed seal knocking vocalization. The spectrogram was computed using a 1024-point Hann window with 87.5% overlap and a 1024-point discrete Fourier transform.

### Data preprocessing

Prior to analysis, the annotated knocking recordings were downsampled from 96 kHz to 19.2 kHz. As knocking calls are low-frequency, this substantially reduced computational cost while retaining the full frequency range relevant to the call.

### Pulse repetition rate estimation

The pulse repetition rate (PRR) of a call is the frequency at which its component pulses are emitted, typically measured in pulses per second (Hz). Saimaa ringed seal knocking calls are characterized by periodic waveforms that repeat at a constant fundamental frequency (Fig. 2), a property closely paralleled in human speech, where pulses repeat at a fundamental frequency (Gerhard 2003). This parallel motivated testing classical fundamental frequency estimation methods, well established in speech-processing research, as a means of automatically estimating the PRR of knocking calls.

To identify which of these methods best captured PRR in knocking calls, we implemented and benchmarked five widely used estimators, spanning both time-domain and time-frequency-domain approaches: the autocorrelation function (ACF; Rabiner 1977), the average magnitude difference function (AMDF; Ross et al. 1974), and the YIN fundamental frequency estimator (de Cheveigné & Kawahara 2002), alongside two time-frequency-domain methods that instead compute periodicity from the amplitude envelope of a call’s spectrogram, using (1) autocorrelation and (2) a Fourier transform to find the frequency of highest magnitude. In both time-frequency methods, spectrograms were processed using per-channel energy normalization (PCEN; Wang et al. 2017) to reduce the effect of background noise and highlight transient events. Compared to the time-domain methods, these approaches mitigate the influence of other frequency bands that might obscure the PRR, at the cost of greater computational demand.

Each estimator required tuning to suit Saimaa ringed seal knocking calls specifically. All methods required an expected upper and lower bound for PRR values, which we set at 5--70 Hz based on values reported in previous manual characterizations of the call (Rautio et al. 2009; Mizuguchi et al. 2016; Young et al. 2025); further testing confirmed these bounds were appropriate. YIN was implemented using the efficient method of Guyot (2018), with a harmonic threshold of 0.7 set through experimental tuning. For the time-frequency-domain methods, spectrograms were computed using a 1024-point discrete Fourier transform (DFT) with a 256-sample Hann window and 78.5% overlap, values selected by testing different parameter combinations and visually assessing how well they separated knocking events from background noise. To validate results from the automatic methods, estimates were compared with manual characterizations of a subset of 200 randomly selected events for which pulse repetition rate could be manually measured. This subset was chosen to provide a tractable evaluation sample while aiming to capture the variability of the full dataset. The mean absolute error (MAE) and its standard deviation (SD) were computed as performance metrics.

### Classification data

The classification dataset consisted of equal-length audio segments with fixed durations of 1.3 seconds, a length chosen to represent the typical duration of a knocking call. One segment was extracted from each annotated knocking event to form the positive (knocking) samples. To simulate the variability introduced when applying a fixed-length analysis window to a continuous real-world audio stream, the 1.3-second window was not aligned with the annotated event boundaries. Instead, its position was randomly shifted within the annotation interval to capture different portions of the event. Negative (noise) samples were obtained in the same manner from all of the available non-knocking events and from randomly selected unannotated segments of background noise. The number of background noise samples, comprising 94.16% of the noise dataset, was selected to ensure that the total amount of negative instances matched the number of positive samples.

The resulting knocking and noise dataset was then converted into three different representations, each suited to a different type of classification model. In all cases, each sample first had its mean subtracted to remove the baseline (DC) offset. A first set of feature vectors was obtained by applying the PRR estimators to each sample. Along with 11 values produced by the collection of methods, the power and zero-crossing rate of each waveform were included, as they are commonly used in fundamental frequency estimation (see Appendix 1 for descriptions of each feature). Furthermore, the raw waveforms were also used as feature vectors with no further processing beyond the DC offset removal. Finally, spectrograms were computed using a 1024-point DFT with a 256-sample Hann window and 78.5% overlap, matching the settings of the time-frequency-based PRR estimation methods.

### Classification models and evaluation

Three classifier types were tested, matched to one of the input representations above. The models used were a support vector machine (SVM; Hearst et al. 1998) for PRR estimation features, a multilayer perceptron (MLP; Hornik et al. 1989) for waveforms, and a convolutional neural network (CNN; LeCun & Bengio 1998) for spectrograms. Comparing these three approaches allowed us to test whether the pulse-rate information estimated earlier in this study is itself sufficient for reliable detection, or whether models that learn directly from the raw acoustic signal or its spectrogram perform better. The MLP and CNN architectures (Table A2 and Fig. 3, respectively) were designed to balance classification performance and computational efficiency, but no thorough optimizations were performed.

**Fig 3.**
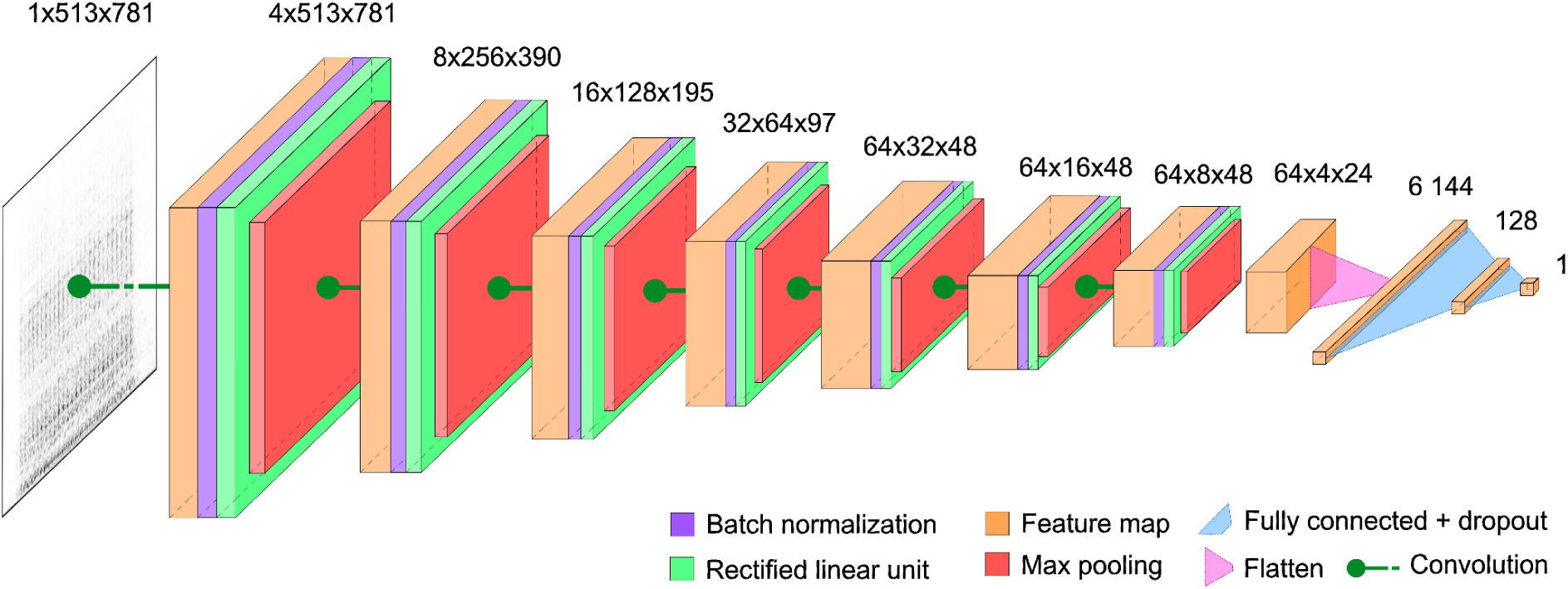
Architecture scheme of our convolutional neural network spectrogram classifier. The dimensions of the input spectrogram and the initial feature maps correspond to (number of channels x height x width).

To evaluate the classifiers across the entire dataset, we used nested cross-validation (nested CV; Varma & Simon 2006) with five outer folds and three inner folds. Compared with conventional CV or evaluation using a single held-out test set, nested CV provides an estimate of generalization performance that accounts for variability across data partitions while reducing optimistic bias arising from hyperparameter selection. Our nested CV implementation consisted of dividing the data into five equal-sized, random, non-overlapping folds, iteratively training the model on four of them (80% of the data), and evaluating it on the remaining, unseen outer fold. To select the optimal hyperparameter combination, each subset of 80% of the data was further divided into three inner folds to optimize the hyperparameters. Two of those inner folds (66% of the subset) were used to train the models with all hyperparameter configurations, and the remaining inner fold (33% of the subset) to evaluate them.

Then, the best-performing combination was used to train the model on the four outer folds. The hyperparameter space was explored using a grid-search approach, in which all values were tested against each other in all possible combinations (Table A3). Results were presented as the mean F1-score across the five outer folds of unseen data, with the corresponding standard deviation.

While nested CV provides an estimate of model performance when data from all four hydrophones are considered jointly, it does not assess the model’s ability to generalize to unseen data e.g. from different geographical locations or deployment periods. Therefore, as a complementary analysis, we evaluated the generalizability of the best-performing classifier type by first aggregating data from three hydrophones to train the classifier and tune its hyperparameters, and then using completely unseen data from the remaining, held-out hydrophone to measure its performance. We did this iteratively for all four hydrophones. In addition, to obtain results more robust to the variability inherent in training a model and selecting a subset of data for hyperparameter optimization, we repeated each experiment five times and reported the mean and standard deviation of the performance metrics over runs. Because H_D_ was deployed in a different breeding season (2022) from H_A,_ H_B,_ and H_C_ (2023), this analysis reflects both spatial and inter-annual variation.

Furthermore, we also assessed whether each model could realistically keep pace with the volume of data PAM generates, which is central to its usefulness as a monitoring tool. This was measured using the real-time factor (RTF), which is defined as the time a model takes to process a waveform and classify it, divided by the duration of the sample. An RTF below one indicates a model can process audio faster than it plays back, meaning it could, in principle, keep up with continuously accumulating recordings. We calculated the RTF over 100 randomly selected samples for each classifier and reported the mean values. Since RTF is inherently hardware-dependent, we report the specific resources used for each model to support reproducibility and fair comparison. To conduct this experiment, we ran our SVM on an Intel Xeon E5-2630 v4 CPU running at 2.2 GHz, and used an NVIDIA GeForce GTX TITAN X GPU with 12 GB of memory for our MLP and CNN. While direct comparison between the SVM and the deep learning-based models is not possible due to differing hardware, using a GPU to accelerate the MLP and CNN reflects standard practice, as such resource-intensive models benefit from the parallel-processing capabilities of GPUs.

### Code and data availability

All code used in our analyses is available at https://github.com/AdriaSolana/ringed-seal-knocking-detection. The data underlying this project will be made available upon request from reviewers or editors during the review process, and publicly deposited on Zenodo upon acceptance of the manuscript.

## Results

### Pulse repetition rate estimation

We ran 200 randomly selected knocking calls through our five PRR estimators, compared their estimates against manual evaluations, and reported their MAE and corresponding SD (Table 1). We then used the best-performing method to estimate the PRR distribution of the knocking calls across hydrophones (Fig. 4).

**Fig 4.**
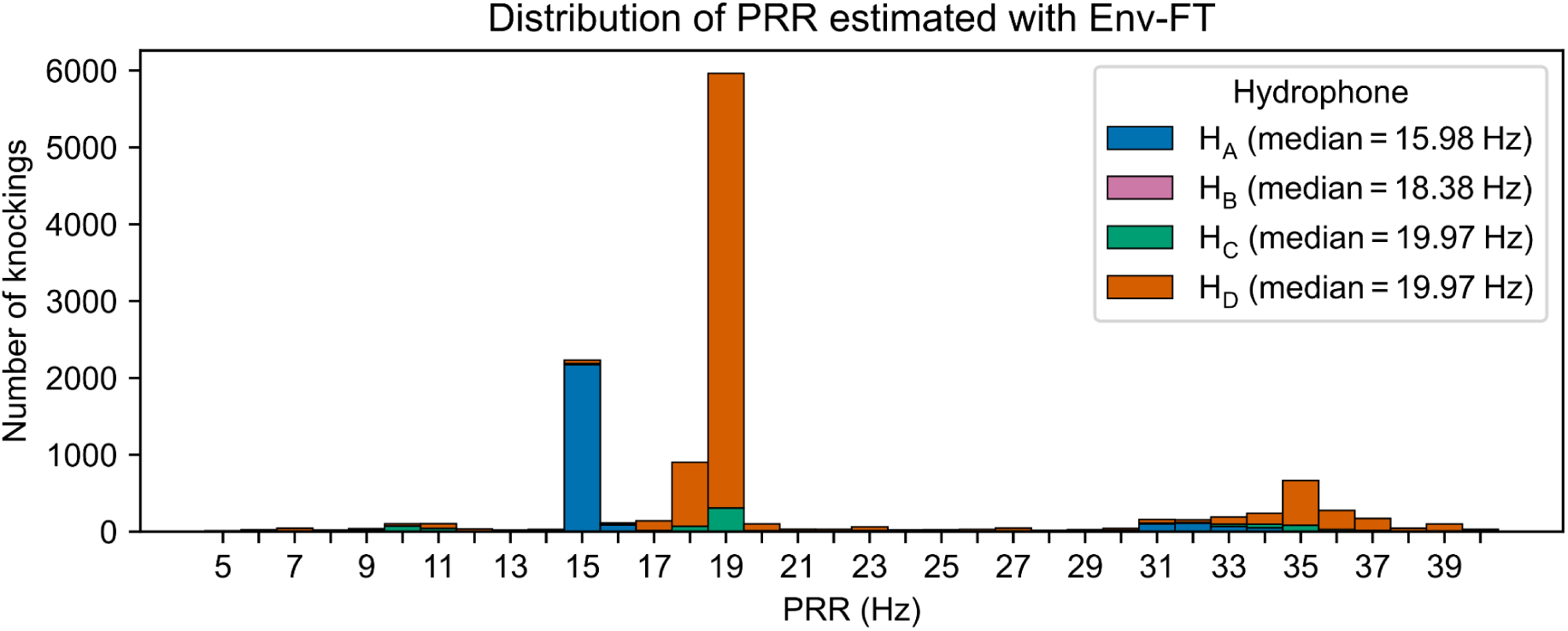
Distribution of Pulse Repetition Rates (PRRs) estimated with Env-FT. The plot was limited to 40 Hz due to the lack of estimates in the higher range.

**Table 1.**
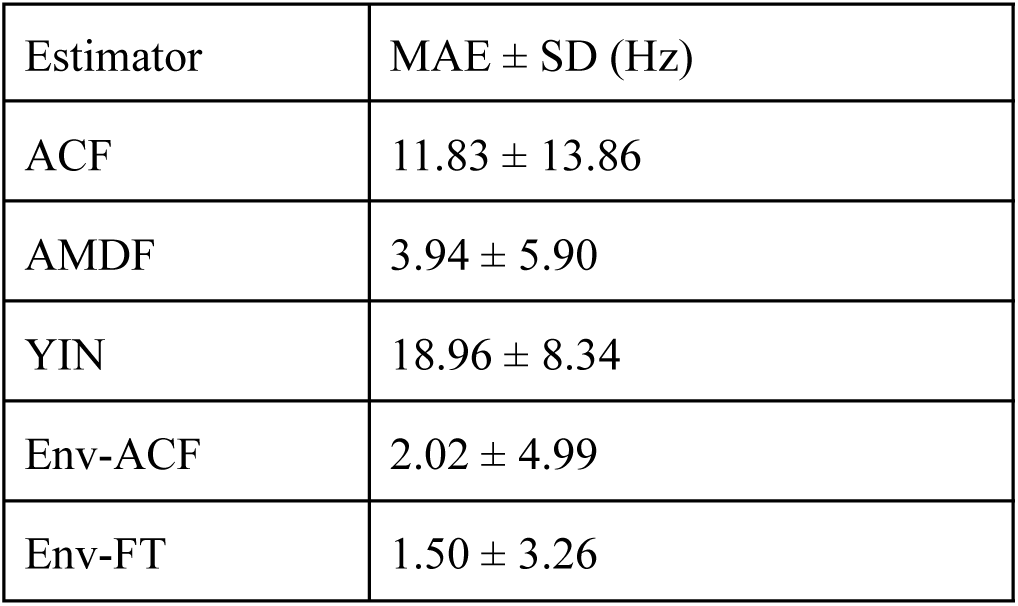
Mean absolute error (MAE) ± standard deviation (SD), in Hz, of 200 automated pulse repetition rate (PRR) estimates relative to the ground-truth PRR. Abbreviations: ACF, autocorrelation function; AMDF, average magnitude difference function; YIN, YIN fundamental frequency estimator; Env-ACF, envelope-based autocorrelation function; Env-FT, envelope-based Fourier transform.

| Estimator | MAE ± SD (Hz) |
| --- | --- |
| ACF | 11.83 ± 13.86 |
| AMDF | 3.94 ± 5.90 |
| YIN | 18.96 ± 8.34 |
| Env-ACF | 2.02 ± 4.99 |
| Env-FT | 1.50 ± 3.26 |

The results exhibited a large variability across methods, with the spectrogram-based estimators performing better. In particular, Env-FT achieved the best results with the lowest MAE and standard deviation. Subsequently, we estimated the PRR distribution over the four hydrophones with Env-FT (Fig. 4). The median PRR was broadly similar across three of the four hydrophones (H_B_: 18.38 Hz; H_C_ and H_D_: 19.97 Hz), while H_A_ showed a lower median (15.98 Hz).

### Classification

Nested CV was used on the three classifiers to obtain robust estimates of performance on unseen data, and, in addition, RTFs were computed over 100 random samples. The classifiers achieved varying F1-scores across the five outer folds, whereas all three models processed audio faster than real-time (RTF < 1), with the CNN and MLP doing so substantially faster than the SVM (Table 2).

**Table 2.** Classifier results from nested cross-validation and real-time factor (RTF) computation. Abbreviations: SVM, support vector machine; MLP, multi-layer perceptron; CNN, convolutional neural network; SD, standard deviation.

| Classifier | Input features | Metrics (mean ± SD) | | | |
| --- | --- | --- | --- | --- | --- |
|  |  | Precision % | Sensitivity % | F1-score % | RTF |
| SVM | PRR features | 98.22 ± 0.33 | 97.18 ± 0.24 | 97.70 ± 0.20 | 0.779 ± 0.253 |
| MLP | Waveforms | 40.00 ± 37.42 | 40.02 ± 48.97 | 26.71 ± 32.62 | 0.001 ± 0.001 |
| CNN | Spectrograms | 99.34 ± 0.11 | 99.22 ± 0.20 | 99.28 ± 0.07 | 0.015 ± 0.008 |

Having achieved the highest and most consistent F1-score, and analyzing events at a speed that would allow for real-time computation, we selected the CNN as our overall best-performing model. Subsequently, we used this classifier for the following experiment, consisting of training the model and optimizing its hyperparameters on a dataset comprised of data from three hydrophones, and evaluating its generalization capabilities on completely unseen data from the remaining, held-out hydrophone. Results show slightly lower F1-scores than those of the nested CV experiments, while being indicative of a high generalization capability across hydrophones (Table 3). Note that the held-out H_B_ dataset comprised only 48 test samples, substantially fewer than the other three hydrophones, which limits the robustness of the performance estimate.

**Table 3.** Results from the best-performing CNN trained on three hydrophones and evaluated on the remaining, held-out hydrophone over five runs. The number (No.) of test samples is the sum of knocking and noise events from each dataset. Abbreviations: hydro, hydrophone; FNR, false negative rate; SD, standard deviation.

| Held-out hydro. | No. test samples | Metrics (mean ± SD) % | | | |
| --- | --- | --- | --- | --- | --- |
|  |  | Precision % | Sensitivity % | FNR % | F1-score % |
| H<sub>A</sub> | 5 242 | 98.85 ± 0.16 | 94.21 ± 1.08 | 5.79 ± 1.08 | 96.47 ± 0.63 |
| H<sub>B</sub> | 48 | 96.00 ± 0.00 | 100.00 ± 0.00 | 0.00 ± 0.00 | 97.96 ± 0.00 |
| H<sub>C</sub> | 1 906 | 99.32 ± 0.35 | 86.40 ± 11.57 | 13.60 ± 11.57 | 91.97 ± 6.69 |
| H<sub>D</sub> | 17 934 | 99.16 ± 0.15 | 97.33 ± 0.45 | 2.67 ± 0.45 | 98.24 ± 0.23 |

## Discussion

Our results showed that both the fine-scale structure and the presence of Saimaa ringed seal knocking calls can be detected automatically, and with high accuracy, from PAM recordings. This demonstrates that automated approaches are capable of substantially supporting the analysis of large passive acoustic monitoring datasets for the Saimaa ringed seal.

### Automated detection and characterization of knocking calls

The five studied pulse repetition rate estimators presented varying performances, with the two time-frequency-domain methods producing more accurate estimates against a ground truth of 200 manually analyzed knocking calls. Env-FT achieved the best results, and its automated estimates agreed closely with an independent PRR estimate derived from the call parameters reported in Young et al. (2025), even though that estimate had been calculated from data collected for a different purpose. This agreement, together with the method’s low error against manual measurement, supports Env-FT as a reliable means of automatically characterizing knocking call structure.

Regarding classification, the separability inherent in the data allowed for various data representations and architectures to yield satisfactory results. The worst-performing strategy was to train a multilayer perceptron on waveforms. An analysis of the training metrics revealed that the MLP was not able to generalize to unseen samples due to its large size compared to the variability of the data; the model complexity was constrained by the size of the input features, namely waveforms with 24 960 samples, and this architecture was not optimal for the problem. In general, MLPs lack temporal invariance and are not a practical solution for such end-to-end systems. There are alternative architectures designed for raw waveform classification in the domain of speech analysis that could outperform our basic MLP, including SincNet (Ravanelli & Bengio 2018) and RawNet2 (Jung et al. 2020). Moreover, both the SVM (using PRR-derived features) and the CNN (using spectrograms) performed considerably better than the MLP. While the SVM is lighter and easier to interpret, its PRR feature extraction increased its RTF, and its F1-score was slightly lower than the CNN’s. The use of spectrograms and convolutional architectures is consistent with recent advances in deep-learning-based acoustic detection of seals and other marine mammals (e.g., Escobar-Amado et al. 2022; Licciardi & Carbone 2024; Tyshko et al. 2024), and the CNN provided the strongest overall balance between classification performance and computational efficiency. A potential improvement to our CNN could consist of adding a Res2Net block (Gao et al. 2021), as evaluated for speech classification tasks by Zhou et al. (2021), to enable multi-scale analysis without generating additional spectrograms.

Further experiments showed that our CNN trained on all but one hydrophone performed remarkably well when evaluated on the held-out hydrophone. This is consistent across the four hydrophones, achieving a minimum F1-score of 91.97% on H_C_. Because H_D_ was recorded in a different breeding season from the other hydrophones, this result reflects generalization across both site and deployment period. Interestingly, performance did not drop on H_A_ (F1-score = 96.47%) despite some variation in call structure (PRR), suggesting that the model is robust at least to this degree of acoustic variation. However, achieving a low false negative rate remained a challenge consistently across all four data partitions, indicating that the model was overly conservative in classifying samples as knocking calls. Given the cost of missing potential seal activity due to the species’ endangered status, future work should focus on improving sensitivity without producing an excessive number of false positives. Overall, these results show that the model is capable of capturing the variability present in the calls and generalizing reasonably well beyond its training conditions.

The CNN’s exceptionally strong performance likely reflects, at least in part, the stereotyped nature of the knocking call itself. The repetitive pulse structure produces a distinctive spectrographic pattern readily distinguishable from most background sounds. This may make knocking vocalizations particularly amenable to automated detection. Despite this high performance, practical deployment requires consideration of false detections, as even low false-positive rates can generate substantial numbers of candidate detections when processing the large volumes of data produced by long-term monitoring. Automated detection should therefore currently be viewed as a tool for substantially reducing, rather than eliminating, the need for manual review, with a workflow of automated screening followed by targeted human verification likely the most effective approach for large monitoring datasets.

### Implications for Saimaa ringed seal conservation

Previous work established knocking vocalizations as a biologically meaningful indicator of breeding-season behavior in the Saimaa ringed seal and identified passive acoustic monitoring as a valuable tool for studying this otherwise inaccessible period of the species’ life cycle, while noting manual analysis as a key constraint on scaling this approach further (Young et al. 2025). The present results speak to this constraint directly. The developed CNN was capable of detecting knocking calls with a mean F1-score of 99.28%, close to the reliability expected of manual review. Importantly, it was able to process audio substantially faster than real time (RTF = 0.015). Furthermore, it maintained high performance (F1-score > 91.97 %) when deployed on data with different characteristics (PRR), from different time periods and geographical locations. These results suggest that automated detection could substantially reduce the effort currently required to analyze acoustic recordings from this population. Given the established link between vocal activity and reproductive behavior (Young et al. 2025), the ability to process PAM data relatively quickly and easily points toward a potential future route for tracking a reproductively critical period of the species’ annual cycle at a scale not previously achievable.

Automated PRR estimation offers a further, complementary contribution by capturing a fine-scale property of the call itself. In other species, variation in properties like this has been linked to factors such as identity, sex, or condition (Manser 2016), raising the possibility that call structure could carry similar information in the Saimaa ringed seal. Automating the measurement of these characteristics makes it feasible to examine questions about calling behavior at the population scale that manual review alone could not practically support. The lower median PRR observed at one hydrophone in this study illustrates the kind of pattern this method can reveal. While investigating the potential mechanisms behind this call structure variation was beyond the scope of the present study, it offers potential avenues for future research. Combined with the detection capabilities described above, this points toward passive acoustic monitoring becoming an increasingly capable tool for understanding and monitoring the Saimaa ringed seal.

### Conclusions

This study developed and evaluated automated methods for detecting and characterizing Saimaa ringed seal knocking vocalizations, addressing the manual-analysis bottleneck that currently limits the scale at which PAM can be applied to this population. Our results indicate that both detection and fine-scale characterization of the species’ only consistently documented call type can be achieved automatically, at a level of accuracy that approaches manual review. While this study explores to what extent these results can be extrapolated across a limited number of sites, years, and acoustic conditions, further experiments with a greater diversity of data sources should be conducted before these methods can be considered ready for routine operational deployment. Nonetheless, the methods presented here show considerable potential to reduce the manual effort associated with acoustic monitoring and could open new opportunities to investigate vocal activity, strengthening the role of passive acoustic monitoring as a conservation tool for this endangered freshwater seal.

## Acknowledgements

The authors wish to thank Metsähallitus and Miina Auttila, Marja Niemi, and Riikka Alakoski, who contributed to the deployment and collection of the hydrophones, as well as our collaborators at Turku University of Applied Sciences. MY is funded by the Finnish Cultural Foundation [55241449]. This work is also supported by WWF Finland [2421/02.07.02/2010] and Tampereen Särkänniemi Ltd. [351/02.01/2022]. This study is part of the SealHabitat cross-border project that is funded by the EU programme Interreg Aurora.

## Appendices

### Appendix 1. Pulse repetition rate-derived features definition

The following list provides descriptions of the features based on the pulse repetition rate (PRR) used in classification with a support vector machine (SVM).

Power (dB): Overall energy of the signal in decibels.

ZCR: Zero-crossing rate, defined as the rate at which the signal waveform crosses zero. ACF PRR (Hz): PRR estimated with the autocorrelation function method (ACF).

ACF rel. amp.: Amplitude of the maximum value of the autocorrelation curve relative to the energy of the signal (relative amplitude).

AMDF PRR (Hz): PRR estimated with the average magnitude difference function (AMDF) method.

AMDF rel. amp.: Amplitude of the minimum value of the AMDF curve relative to the maximum of the curve (relative amplitude).

YIN PRR (Hz): PRR estimated with a custom implementation of the YIN pitch tracker (YIN) based on Guyot (2018).

YIN harmonic rate: Confidence score of the PRR estimate using YIN.

YIN argmin (Hz): PRR estimated with a custom implementation of YIN, but ignoring the harmonic rate threshold.

PCEN env. ACF (Hz): PRR estimate using the autocorrelation function over an amplitude spectral envelope of a per-channel energy normalization-processed (PCEN) spectrogram.

PCEN env. ACF rel. amp.: Amplitude of the maximum value of the autocorrelation curve relative to the energy of the signal. The curve is obtained with ACF over an amplitude spectral envelope of a PCEN-processed spectrogram.

PCEN env. FT argmax (Hz): Maximum of the Fourier transform of an amplitude spectral envelope of a PCEN-processed spectrogram.

PCEN env. FT entropy: One minus the entropy of the Fourier transform of an amplitude spectral envelope of a PCEN-processed spectrogram.

### Appendix 2. Multilayer perceptron model architecture

**Table A2.** Description of the architecture of our MLP. The abbreviation “ReLU” corresponds to “rectified linear unit”.

| Layer | Type | Input shape | Output shape | Activation |
| --- | --- | --- | --- | --- |
| fc1 | Linear (fully connected) | 24 960 | 64 | ReLU |
| dropout1 | Dropout | 64 | 64 | - |
| fc2 | Linear (fully connected) | 64 | 16 | ReLU |
| dropout2 | Dropout | 16 | 16 | - |
| fc3 | Linear (fully connected) | 16 | 1 | Sigmoid |

### Appendix 3. Explored hyperparameters for hyperparameter tuning

**Table A3.** Hyperparameters tested in nested cross-validation and CNN training. Abbreviations: SVM, support vector machine; MLP, multi-layer perceptron; CNN, convolutional neural network; rbf, radial basis function.

| SVM |  | MLP & CNN |  |
| --- | --- | --- | --- |
| Hyperparameter | Values | Hyperparameter | Values |
| C | 0.1, 1.0, 10 | Batch size | 64, 128, 256 |
| Kernel | linear, rbf | Learning rate | 1e-4, 1e-5, 1e-6 |
| $\gamma$ | scale, 0.01, 0.001 | Dropout | 0.1, 0.5, 0.7 |

